# RNA-sequencing of human post-mortem hypothalamus and nucleus accumbens identifies expression profiles associated with obesity

**DOI:** 10.1101/2022.01.08.473382

**Authors:** Christian Wake, Julie A. Schneider, Thor D. Stein, Joli Bregu, Adam Labadorf, Ann McKee, Philip L. De Jager, David A. Bennett, Sudha Seshadri, Richard H. Myers, Anita L. DeStefano

## Abstract

Obesity, the accumulation of body fat to excess, may cause serious negative health effects, including increased risk of heart disease, type 2 diabetes, stroke and certain cancers. The biology of obesity is complex and not well understood, involving both environmental and genetic factors and affecting metabolic and endocrine mechanisms in tissues of the gut, adipose, and brain. Previous RNA sequencing studies have identified transcripts associated with obesity and body mass index in blood and fat, often using animal models, but RNA sequencing studies in human brain tissue related to obesity have not been previously undertaken. We conducted both large and small RNA sequencing of hypothalamus (207 samples) and nucleus accumbens (276 samples) from individuals defined as consistently obese (124 samples), consistently normal weight as controls (148 samples) or selected without respect to BMI and falling within neither case nor control definition (211 samples), based on longitudinal BMI measures. The samples were provided by three cohort studies with brain donation programs; the Framingham Heart Study (FHS), the Religious Orders Study (ROS) and the Rush Memory and Aging Project (MAP). For each brain region and large/small RNA sequencing set, differential expression of obesity, BMI, brain region and sex was performed. Analyses were done transcriptome-wide as well as with a priori defined sets of obesity or BMI-associated mRNAs and microRNAs (miRNAs). There are sixteen mRNAs and five microRNAs that are differentially expressed (adjusted p < 0.05) by obesity or BMI in these tissues, several of which were validated with qPCR data. The results include many that are BMI-associated, such as APOBR and CES1, as well as many associated with the immune system and some with addiction, such as the gene sets “cytokine signaling in immune system” and “opioid signaling”. In spite of the relatively large number of samples, our study was likely under-powered to detect other transcripts or miRNA with relevant but smaller effects.

## Introduction

Obesity is the accumulation of body fat to the point of excess that may cause serious negative health effects. These health risks are all among the world’s leading causes of death and include heart disease, type 2 diabetes, stroke and certain cancers (breast, colon, kidney, endometrial, gallbladder, and liver). The prevalence of adult obesity in developed countries has increased dramatically over time until leveling off in the last decade at approximately 35% [1] but continues to increase worldwide. [2]

The etiology of obesity is complex, influenced by genetic and environmental factors such as diet, caloric intake and amount of physical activity. Many genome-wide association studies (GWAS) [3–14] have been conducted in order to find specific sequence variants associated with obesity, and at least 75 genetic loci have been identified [15, 16]. Although the functional consequences of these variants are not all understood, many implicated genes have functions associated with the neuro-endocrine and related neuronal systems. These include FTO (fat mass and obesity associated gene), the strongest and first association made to BMI via GWAS, implicated through a cluster of BMI-associated SNPs in its first intron. FTO is expressed ubiquitously but is especially highly expressed in the brain and in neurons, where it may act in the sensing of intracellular amino acid concentrations [15, 17]. Other neuroendocrine genes among the GWA-implicated set are MC4R (melanocortin receptor 4), NPC1 (NPC Intracellular Cholesterol Transporter 1), NRXN3 (Neurexin3) and (Potassium Calcium-Activated Channel Subfamily M Alpha 1). MC4R is a hormone receptor in the brain regulating the melanocortin system on which the hunger hormones leptin and ghrelin act, NPC1 is involved in membrane cholesterol transport in glial cells, NRXN3 is a neuronal cell surface protein associated with cell adhesion, and KCNMA1 is key in neuron excitability.

Neuroendocrine involvement in the development and subsequent effects of obesity is unsurprising, given the association of obesity with dysregulation of energy homeostasis, and the relevance of neuroendocrine function to hunger, diet, and response to the metabolic hormones such as insulin and leptin. Consequently, we were motivated to interrogate obesity via RNA sequencing in the brain. We chose to study the hypothalamus, as this brain region is heavily involved in regulating the endocrine system. The hormones leptin and ghrelin act in the hypothalamus, increasing or decreasing satiation after being produced in the adipose tissue or gut, respectively [18]. In addition to the hypothalamus, we conducted RNA sequencing in the nucleus accumbens. This brain region plays a significant role in reward systems, pleasure, and impulse responses, which have been implicated in obesity related behaviors [19]. While brain related metabolic, endocrine and behavioral effects are likely to contribute to obesity, these have not been previously evaluated in human brain.

Measurements of gene expression are a key tool that that may assist in identifying obesity associated genes and the underlying variants responsible for the relationship. RNA sequencing yields transcript abundance estimates which may be affected by all of the factors that GWAS can capture and those it cannot, including genetic, environmental and gene regulatory effects. Furthermore, RNA sequencing analysis may offer a more direct measure of the functional state of a cell or tissue, showing which transcripts are being utilized and may be particularly useful for interrogating the effect of disease on or within specific tissues. RNA expression analyses of obesity and BMI have been conducted in human adipose [2, 20–25] and in animal models [26, 27], but not within human brain tissue. RNA sequencing analysis contrasting post-mortem human brain tissue of obese and normal weight individuals has the potential to reveal novel significant insights into the mechanisms of obesity.

We have conducted large and small RNA sequencing on human hypothalamus and nucleus accumbens samples from The Framingham Heart Study, The Religious Orders Study and the Memory and Aging Project. Differential expression analyses of obese versus controls were conducted with individuals selected based on strict longitudinal BMI requirements, and analyses of continuous BMI done with a larger set of samples. Here we present transcriptome-wide and obesity-implicated gene and microRNA differential expression results of obesity and BMI, obesity gene set enrichment results, and similar analyses of differential expression by brain region and by sex with the same data.

## Materials and Methods

### Sample Information

Approximately 100 mg of post-mortem frozen hypothalamus and nucleus accumbens tissue were provided by the brain banks at three cohort studies; the Framingham Heart Study (FHS), Framingham, Massachusetts, and the Religious Orders Study (ROS) and the Memory and Aging Project (MAP) centered at the Rush University Medical Center, Chicago, Illinois. The FHS brain donation program represents only a portion of all FHS participants as it was established after the study initiation and not all participants chose to consent to brain donation. Brain donation is part of the ROS and MAP study protocols and each participant signed an Anatomic Gift Act in addition to informed consent and a repository consent allowing their biospecimens and data to be repurposed. Details of study design for each of these cohorts has been previously published [28–30]. Dissection of frozen hypothalamus and nucleus accumbens was done by a single pathologist at each site. FHS, ROS and MAP each collect detailed phenotypic, genotypic and neuropathological data, including longitudinal measure of BMI, enabling application of strict criteria sample selection. Cases were defined as individuals who were consistently obese (BMI>30) for at least 4 consecutive measures prior to death. Controls were defined as individuals who were consistently normal weight (18< BMI < 25) for at least 4 consecutive measures prior to death. Over cases and controls this represents a time span ranging from 2.6 years to 23.9 years, with a median span of 3.2 years. The initial phase of sample selection and sequencing consisted of 230 cases and controls, followed by a second phase of 291 samples, including 32 cases and 25 controls and the remainder a population sample that did not include individuals meeting the case or control definitions but were otherwise not selected with respect to BMI.

### Sample Preparation and Sequencing

Total RNA from these samples were isolated using QIAzol Lysis Reagent (Qiagen, Valencia CA) and purified using miRNeasy MinElute Cleanup columns. RNA integrity was evaluated with per-sample RNA Integrity Number (RIN) using Agilent’s Bioanalyzer 2100 and RNA 6000 Nano Kits.

Large RNA library preparation and sequencing was conducted at the Oklahoma Medical Research Foundation Genomics Core using the Illumina TruSeq Stranded Total RNA with Ribo-Zero Gold (rRNA-depleted) preparation kits. The sequencing was conducted in ten separate batches (sequencing runs conducted on different flow cell and/or day), four in phase 1 and six in phase 2. In order to eliminate lane effects within each batch, samples were barcoded to distinguish samples and multiplexed across lanes. Sequencing of 2×150 paired-end reads was done using the Illumina Hiseq 3000 with target read depth of 40 million read pairs.

Small RNA library preparation and sequencing at the University of California, Los Angeles, Clinical Microarray Core using New England Biolabs NEBNext library preparation, and sequencing of 1×50 single-end reads done on the Illumina HiSeq 3000 with target read depth of 15 million. Sequencing was conducted in twenty-seven batches, eleven in phase 1 and sixteen in phase 2, and samples were multiplexed within each batch to eliminate any lane artifacts.

### Sequencing Analysis

Sequencing reads were clipped of adaptors and length-filtered using the tool cutadapt v 1.14 [31]. Clipped large RNA sequencing reads shorter than 50 nucleotides were removed, and clipped small RNA sequencing reads shorter than 15 or longer than 23 were removed. For quality control, the nucleotide trimming tool sickle v 1.33 was applied with Phred quality threshold of 20, removing low quality ends of reads and once again applying the lower bound read length filter [32]. Reads of each sequencing sample were aligned to the hg38 human reference genome with the alignment tool STAR, Spliced Transcript Alignment to a Reference, v 2.5.3a [33]. Large RNA alignments with mismatches more than 5% of the read length and small RNA alignments with greater than 1 mismatch were disallowed. Spliced reads were disallowed for the small RNA sequencing samples. Read counting of small RNA alignment files was done using the HTSeq intersection non-empty method [34] implemented in Verse v. 0.1.5 [35], and miRBase version 22 [36] mature microRNAs. Large RNA abundance estimation was done using the method RSEM [37] with input STAR alignment of reads to gencode version 23 genes translated into transcriptome coordinates (STAR’S –quantMode TranscriptomeSam option).

### Sample Sets analyzed

Sample filtering based on RIN and Braak neurofibrillary tangle stage [38], a measure of neurodegeneration, was applied yielding three separate analyses defined by different filter criteria. In all analyses, 39 samples with RIN less than 3 were removed. Our choice of RIN > 3 as a lower threshold was guided by observable differences at very low RIN values. Samples with RIN less than 3 have much lower read depth and more multi-mapped reads than samples above 3. The three filtering criteria used are notated as RIN6/BraakIV, RIN3/BraakIV and RIN3/Braak-adjusted and are described in detail below. The sample sizes of obesity and BMI differential expression analyses with varying sample filters (both sexes combined) are shown in Table 1.

**Table 1:**
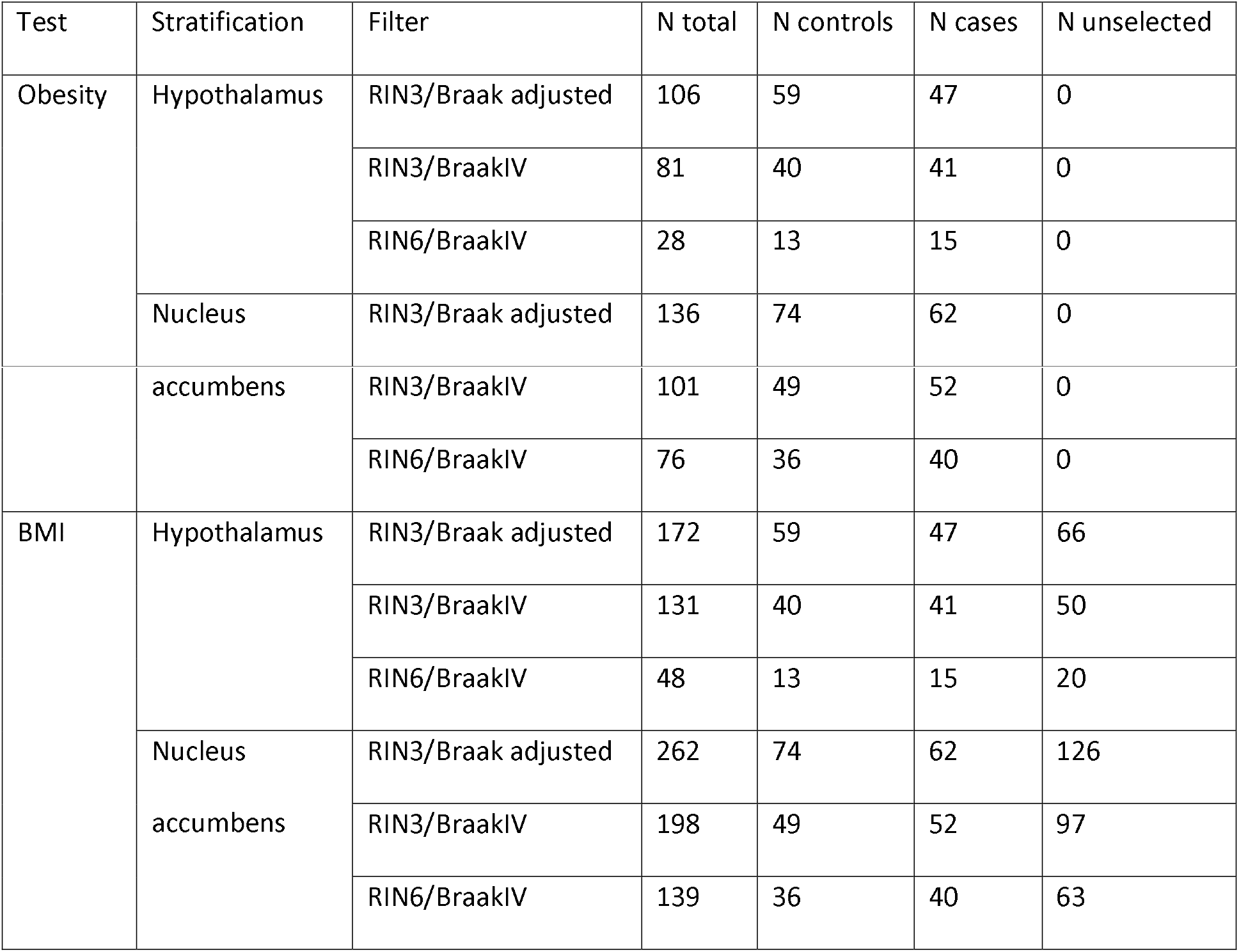
Sample sizes of the three RIN and Braak stage filters from the obesity and BMI differential expression analyses, without sex stratification.

RIN3/Braak Adjusted: This analysis included samples with RIN >= 3 and no filtering on Braak stage. Although RIN >= 6 is a more common RIN filtering criteria, we chose a lower threshold for our main analyses because of our choice of sequencing library preparation with ribosomal RNA depletion, which is more resilient to degraded RNA than poly-A selection. In addition, we did not observe the differences in quality noted for our RIN < 3 samples for samples with RIN between 3 and 6. Given these factors, we wanted to retain a larger sample size.

Braak stage was accounted for with adjustment in the differential expression models. However, covariate adjustment may not sufficiently account for the effect of substantial neurodegeneration, particularly because neurodegeneration is not randomly occurring across cases and controls. Obesity has been previously associated with neurodegeneration [39], and this is seen in these samples, in which obesity is associated with Braak stage independent of age of death. Although lower age of death is associated with lower Braak stage, obesity is not associated with age of death in this data. In order to mitigate confounding effects from the association of obesity with Braak stage, we have also conducted the RIN3/BraakIV analysis.

RIN3/BraakIV: To more thoroughly address the potential effect of neurodegeneration indicated by a high Braak stage, we conducted an additional analysis with a Braak filter excluding subjects with Braak stage greater than IV. As with all analyses, only samples with RIN >= 3 were retained. The Braak filter reduced the number of subjects from 293 to 221 and samples (both brain regions) from 483 to 365. The loss of power by the reduction of sample size by nearly one quarter may be offset by greater homogeneity in the remaining samples after excluding those with Alzheimer’s disease pathology.

RIN6/BraakIV: A third analysis was done excluding subjects with Braak > IV and applying a more stringent and standard RIN filter excluding samples with RIN < 6. Although RIN is also accounted for in the models, this greater homogeneity of RIN values may give us more confidence that any effects of RNA degradation on gene quantifications has not affected the differential expression analysis. However, this final RIN6/BraakIV filter reduces the RIN3/BraakIV sample size to 189, nearly by half. The benefit for increased RIN stringency is likely negated by the loss of power from reduced sample size.

### Differential Expression Analysis

Differential expression analyses were conducted to evaluate expression differences between the obese and normal weight samples and for association with body mass index (BMI). Hypothalamus and nucleus accumbens samples were analyzed separately, and sex stratified and combined analyses were conducted. For each sample subset, genes and microRNAs with zero counts in more than half of samples were excluded from analyses.

Differential expression testing of obesity status was performed with two methods: DESeq2 and linear regression with LIMMA. Normalization of gene abundance estimates and raw miRNA counts was conducted using the DESeq2 normalization method prior to DESeq2 differential expression tests. Normalization was performed using the DESeq2 rlog method (following ComBat batch correction) for LIMMA tests. Differential expression testing of last measured BMI was performed using linear regression with LIMMA. Last BMI was used instead of mean BMI because the two measures are highly correlated in these data and the last BMI was measured more closely to death, after which tissue used for expression analysis was obtained and frozen. Principle Component Analyses showed clear differences by sequencing phase but not sequencing batch, so both LIMMA tests (obesity and BMI) applied ComBat batch correction of phase to DESeq2 rlog transformed data, gene abundance estimates or microRNA raw counts. Each test applied a false discovery rate (FDR) adjusted p-value significance threshold of 0.05.

Phase of sequencing and RIN were included as covariates in the models for all tests, with additional covariates chosen based on association with the model outcome (gene abundance estimates or microRNA counts). Potential covariates were added to the model only if they were significantly associated with more than 10% of the normalized counts across the full set of genes or microRNAs. The tests of association with covariates were done with ANOVA, logistic regression, and Spearman correlation for batch, study, and age of death, respectively. Covariates considered were sex, study of origin (FHS or ROSMAP), age at death, sequencing batch (in which case sequencing phase is removed). The association tests used were logistic regression for sex and study, Spearman correlation for age at death, and ANOVA for sequencing batch. Final covariates used for each sample subset may differ, and are shown in Table 2. For these tests and all differential expression analyses, Bioconductor v. 3.0 and R v. 3.1.1 were used.

**Table 2:** Covariates used for each obesity and BMI differential expression analysis (RIN included in all cases so is not shown in table).

| Test | Sample set | RIN3/Braak<br>adjusted | RIN3/BraakIV | RIN6/BraakIV |
| --- | --- | --- | --- | --- |
| Obesity | Hypo/Male | Phase, Braak | Phase | NA: sample size<br>too small |
|  | Hypo/Female | Phase, Braak,<br>Age_Death, Study | Phase | Phase |
|  | Hypo/BothSexes | Phase, Braak,<br>Age_Death, Study | Phase | Phase |
|  | NucAcc/Male | Phase, Braak,<br>Age_Death | Phase,<br>Age_Death | Phase |
|  | NucAcc/Female | Phase, Braak | Phase | Phase |
|  | NucAcc/BothSexes | Phase, Braak | Phase | Phase |
| BMI | Hypo/Male | Phase, Braak | Phase | Phase |
|  | Hypo/Female | Phase, Braak,<br>Age_Death, Study | Phase,<br>Age_Death | Phase |
|  | Hypo/BothSexes | Braak,<br>Age_Death,<br>Study, Batch, Sex | Phase,<br>Age_Death, Study | Phase |
|  | NucAcc/Male | Braak,<br>Age_Death,<br>Study, Batch | Phase,<br>Age_Death, Study | Phase,<br>Age_Death, Study |
|  | NucAcc/Female | Phase, Braak,<br>Age_Death, Study | Phase,<br>Age_Death | Phase,<br>Age_Death, Study |
|  | NucAcc/BothSexes | Braak,<br>Age_Death,<br>Study, Batch, Sex | Age_Death,<br>Study, Batch, Sex | Age_Death,<br>Study, Batch, Sex |

### Differential expression of obesity implicated mRNA and miRNA

In addition to transcriptome-wide analyses, a focused assessment of mRNAs and miRNAs previously implicated in obesity and obesity-related phenotypes was conducted. An FDR adjustment correcting only for the reduced set of hypotheses was applied to the nominal DESeq2 p-values of the mRNAs and miRNAs in the obesity set.

The mRNA subset was determined using results of a BMI GWAS meta-analysis [40]. Genes whose genomic location overlap LD blocks of the most significant BMI-associated SNPs were included in our subset, for a total of 549 genes. The miRNA subset was taken from Deiuiliis [41] which contains a conglomeration of miRNAs previously implicated with obesity and related traits from studies using a variety of study designs, tissue types and models. The sum of the miRNA set include miRNAs associated with obesity, cardiometabolic disease, adipogensis, lipogenesis, insulin resistance and hepatic glucose homeostatis, totaling 159 of the 2813 miRBase v. 21 mature miRNAs.

### Gene Set Enrichment Analysis

Gene Set Enrichment Analysis (GSEA) [42] was performed with the R package fgsea, Fast Gene Set Enrichment Analysis [43]. The transcriptome-wide mRNA DESeq2 results ordered by the test statistic was used as input. The gene sets from The Molecular Signatures Database (MSigDB) version 6.0 [42, 44], specifically the subset of curated gene sets (C2) and canonical pathways (CP), were tested for significant enrichment based on gene order. The fgsea function was run with 100,000 permutation gene label permutations for p-value estimation (nperm) and filtering pathways with less than 15 or more than 500 genes (minSize, maxSize). This was done for each of the DESeq2 analyses performed. For interpretation of results, gene sets were manually associated with a set of general categories (obesity, neurons, cancer, development, transcription/translation, etc.) based on matching of category key-words within names of gene-sets.

### Validation

Validation of DESeq2 obesity and brain region differential expression results for nine mRNA and miRNA transcripts was performed with the QuantStudio 12K Flex [45] TaqMan Array Card protocol (Life Technologies Corporation, 6055 Sunol Blvd, Pleasanton, CA 94566) using the simultaneous detection of miRNA and mRNA procedure described by Buchholz [46]. Reverse transcription (RT) was performed for 10ng of RNA (5ng/ul concentration). RT was performed by ligating a poly-A tail to the 3’ and an adaptor to the 5’ end of the transcript as described by the manufacturer [47]. mRNA RT was performed using Universal RT primers, random hexamers, and RT enzyme mix according to protocol [46]. The resulting cDNA was then subjected to miRNA pre-amplification for miRNA transcript specific RT amplification using the miR-AMP forward primer, and miR-AMP reverse primers for each miRNA to be assayed according to manufacturer’s protocol [47].

The products were detected by qPCR in the Array Card, using gene-specific mRNA TaqMan Gene Expression Assays and miRNA-specific TaqMan Advanced miRNA Assays which were plated in the Array Card. Each Array Card consists of 384 wells, each containing a specific assays are. Each card has eight slots into which eight individual samples can be applied to 48 wells into which that sample is distributed by centrifugation. Assays were performed in triplicate, permitting up to 15 assays to be assessed within each card for all eight samples, with an Applied Biosystems internal standard in the 16^th^ position.

A total of 200 female samples selected to permit validation of differentially expressed genes and miRNA were assayed for the thirteen target assays. Targets for obesity were USP6, TTN, and NDNF, with a range of mean count values (100 to 3170) and absolute value of log fold-changes (0.73 to 1.78). Targets for brain region were ARPP21, SYNDIG1L, DRD1, hsa-miR-139-5p, hsa-miR-552-5p, and hsa-miR-10b-5p, with a range of mean count values from 130 to 38261 and log fold-change from 1.11 to 6.09. In addition, one mRNA and two miRNA transcripts were assayed (TMEM186, hsa-miR-154-5p and hsa-miR-423-3p), selected from the RNA sequence data to be transcripts with uniform expression across all contrasts, low standard deviations and mean count values representative of the chosen miRNA and mRNA assays.

Analysis of differential expression was performed with a delta-delta-Ct (ΔΔCt) algorithm, as described in Livak and Schmittgen 2001 [48] and implemented with the R package ddCt [49] which yielded ΔΔCt and fold-change values. mRNA and miRNA analyses were done separately, with TMEM186 or hsa-miR-154-5p and hsa-miR-423-3p used as reference transcripts, with TaqMan’s rRNA reference transcript Hs999999-1_s1 used in both cases. Significance of ΔΔCt distribution differences was evaluated by t-test if normality was shown with a Shapiro-Wilk test, or by Wilcoxon rank test if not.

## Results and Discussion

### Large RNA Differential Expression of Obesity and BMI

Significant results from the large and small RNA sequencing DESeq2 and LIMMA differential expression tests of obesity from the RIN3/BraakIV filtering sets are shown in Tables 3 and 4, positive log fold-changes indicating higher expression with higher BMI or obesity status. The large RNA results of obesity and BMI tests from all sets are within Supplementary File 1 (ObesityBMI_Significant_Results.xls), and summarizations of the numbers of results of all obesity, BMI, brain region and sex analyses are shown in Supplementary File 2 (Results_Summary_padj0.05.xls).

**Table 3:** Large RNA differential expression significant results of obesity (DESeq2) and BMI (LIMMA), from RIN3/BraakIV analyses.

| Gene name | Chr | Description | Test | Stratification | LogFC | Adjusted p |
| --- | --- | --- | --- | --- | --- | --- |
| FCGBP | 19 | Fc fragment of IgG binding protein | Obesity | Hypo/M | -1.478 | 0.006 |
| SLAMF8 | 1 | Lymphocyte activation associated | Obesity | Hypo/M | -1.165 | 0.029 |
| HJURP | 2 | Holliday junction recognition | BMI | Hypo/M | -0.447 | 0.044 |
| C1QTNF4 | 11 | Inflammatory regulation, hypothalamic and obesity-associated | Obesity | Hypo/M | 1.078 | 0.049<br>(obesity-set) |
| MEPE | 4 | matrix extracellular phosphoglycoprotein | Obesity | Hypo/F | -3.917 | 3.694E-05 |
| TBR1 | 2 | Cortical development | Obesity | Hypo/F | -3.433 | 0.010 |
| SERPINE1 | 7 | serine proteinase inhibitor family, innate immunity | Obesity | Hypo/F | -1.523 | 0.027 |
| NDNF | 4 | neuron derived neurotrophic factor | Obesity | Hypo/F | 0.990 | 0.027 |
| APOBR | 16 | apolipoprotein B receptor | Obesity | Hypo/Both | -0.545 | 0.007<br>(obesity-set) |
| TTN | 2 | vertebrate striated muscles | Obesity | Hypo/F | 0.725 | 0.025 |
|  |  |  | BMI | Hypo/F | 0.039 | 0.019 |

\*Chr: chromosome, LogFC: log fold-change, Hypo: hypothalamus, NucAcc: nucleus accumbens, M: male, F: female

**Table 4:** Small RNA differential expression significant results of obesity (DESeq2) and BMI (LIMMA), from RIN3/BraakIV analyses.

| ID | Name | Chr | Test | Stratification | LogFC | Adjusted p |
| --- | --- | --- | --- | --- | --- | --- |
| MIMAT0000267 | hsa-miR-210-3p | 11 | Obesity | Hypo/M | -0.493 | 0.003 (obesity-set) |
| MIMAT0000254 | hsa-miR-10b-5p | 2 | Obesity | Hypo/Both | 0.779 | 0.011 (obesity-set) |
| MIMAT0000421 | hsa-miR-122-5p | 18 | Obesity | NucAcc/F | 1.132 | 0.014 (obesity-set) |
| MIMAT0000095 | hsa-miR-96-5p | 7 | Obesity | NucAcc/Both | -0.788 | 0.011 |
| MIMAT0000728 | hsa-miR-375-3p | 2 | Obesity | NucAcc/Both | -1.038 | 0.050 (obesity-set) |

\*Chr: chromosome, LogFC: log fold-change, Hypo: hypothalamus, NucAcc: nucleus accumbens, M: male, F: female

Statistically significant (adjusted p < 0.05) differential expression results for obesity status are sparse for large RNAs in both hypothalamus and nucleus accumbens and both sexes, regardless of the RIN/Braak sample filtering. There are 16 unique large RNAs with significant DESeq2 differential expression between obese and control samples; all from hypothalamus. Two of these are not significant transcriptome-wide but are significant in the obesity-associated set of transcripts (C1QTNF4 and APOBR). There are no mRNA transcripts that are significant in both male and female stratified analyses.

DESeq2 yielded more results than LIMMA. The transcriptome-wide LIMMA analyses of obesity had two significant results: HJURP in hypothalamus/male (filtering RIN3/BraakIV, adjusted p: 0.044, log fold-change: −0.447) and USP6 in hypothalamus/female (filtering RIN6/BraakIV, adjusted p: 0.034, log fold-change: 1.037). Both are also transcriptome-wide significant with DESeq2 in the same sample sets. The LIMMA analyses of BMI yielded only one result, titin (TTN), in the hypothalamus/female from both the filtering RIN3/Braak adjusted (adjusted p: 0.004, log fold-change: 0.0370) and RIN3/BraakIV (adjusted p: 0.019, log fold-change: 0.039), which is also among the DESeq2 obesity results for the hypothalamus/female (filtering RIN3/BraakIV, adjusted p: 0.025, log FC: 0.725).

Several of the obesity and BMI significantly differentially expressed genes are associated with the immune system and vascularization. SLAMF8 is associated with lymphocyte activation and C1QTNF4 with cytokine activity, both significant in hypothalamus, male (filtering RIN3/Braak4). In that sample subset, SLAMF8 has significantly lower expression in obese samples transcriptome-wide (adjusted p: 0.029, log FC: −1.165) and with the obesity-implicated set, C1QTNF4 is significantly more highly expressed in obese samples (adjusted p: 0.049, log FC: 1.078). MUC16 is a member of and FCGBP is associated with the mucin family of proteins which are associated with the immune system and with cancer. Both have significantly lower expression in obese samples relative to control for the hypothalamus, both sexes, filtering RIN3/Braak adjusted (adjusted p: 0.017 and 0.040, log FC: −2.649 and −1.036, respectively) and for FCGBP also in the hypothalamus, male (filtering RIN3/Braak adjusted and RIN3/Braak4, adjusted p: 0.0002 and 0.0055, log FC: −1.653 and −1.478, respectively). CXCL8 has lower expression in obese relative to control samples in the hypothalamus, both sexes (filtering RIN3/Braak adjusted, adjusted p: 0.021, log FC: −1.560) and is associated with inflammation and with angiogenesis. NDNF, associated with revascularization, is more highly expressed in obese samples for hypothalamus, female (filtering RIN3/Braak4, adjusted p: 0.027, log FC: 0.990), and SERPINE1 which inhibits fibrinolysis thus promoting blood clotting, in the same sample set is more lowly expressed in obese samples (filtering RIN3/Braak4, adjusted p: 0.027, log FC: −1.523).

Carboxylesterase 1 (CES1) is significantly differentially expressed by obesity status with DESeq2 in the hypothalamus, both sexes (filtering RIN3/Braak adjusted, adjusted p: 0.035, log fold-change: - 1.287). It is an enzyme that hydrolyzes endogenous and exogenous esters. This includes those of cocaine, heroin and other toxins, but also of cholesteryl esters and triacylglycerols, thus playing a role in both xenobiotic detoxification or activation as well as cholesterol and lipid metabolism. It is abundantly expressed in liver and adipose tissue [50]. Expression and activity of CES1 in adipose tissue has been shown to be increased in obese and type 2 diabetic individuals relative to control [51], although our results show CES1 expression in the hypothalamus is down in obese relative to control samples.

Among our obesity-implicated mRNAs is apolipoprotein B receptor (APOBR), which is significantly differentially expressed with the DESeq2 and LIMMA analyses of obesity in the hypothalamus, both sexes (filtering RIN3/Braak4, adjusted p: 0.007, log FC: −0.545). Its expression is lower in obese relative to control. APOBR is the receptor for APOB, an apolipoprotein produced in the gut and liver that binds lipids to form low density lipoprotein chylomicrons that transport lipids in the blood. The receptor itself binds to APOB, mediating the endocytosis of the lipids of the lipoprotein. A mutation in APOBR has been associated with hypothyroidism [52]. APOBR is known as a macrophage receptor but is known to be expressed in the brain [53], and cholesterol metabolism and transport are often implicated in neurological disease [54]. The large RNA DESeq2 and LIMMA results are shown in Table 3 and Supplemental File 1 (ObesityBMI_Significant_Results.xls), first tab.

### Small RNA Differential Expression of Obesity and BMI

Like the significant large RNA results of obesity and BMI tests, vascularization is again associated with many of the small RNA results. Unlike the large RNAs, the significantly differentially expressed small RNA results are nearly exclusively from the nucleus accumbens rather than hypothalamus. There are only three DESeq2 significant miRNA results from the hypothalamus, but each are among the *a priori* defined obesity-implicated set and each is associated with angiogenesis. Two have lower expression in obese samples relative to control (hsa-miR-210-3p and hsa-miR-375-3p) while one has the opposite expression pattern (hsa-miR-10b-5p). hsa-miR-210-3p is significantly differentially expressed in the hypothalamus, male RIN3/BraakIV (adjusted p: 0.003, log fold-change: −0.493) and RIN3/Braak adjusted (adjusted p: 0.014, log FC: −0.453), hsa-miR-375-3p in hypothalamus, both sexes (RIN3/Braak adjusted, adjusted p: 0.008, log FC: −1. 061), and hsa-miR-10b-5p in the hypothalamus, both sexes (RIN3/BraakIV, adjusted p: 0.011, log FC: 0.779). In addition to angiogenesis, hsa-miR-210-3p is associated with hypoxia, cardiac disease and cancer, and hsa-miR-375-3p with regulation of protein kinase B signaling and endothelial cell apoptosis.

hsa-miR-375-3p is also significant with the DESeq2 test in nucleus accumbens, both sexes (filter RIN3/BraakIV, adjusted p: 0.050, log FC: −1.038), and is the only transcript that is significant in both brain regions in any analysis. There are a number of transcripts from the nucleus accumbens, either female or both sexes (filtering RIN3/Braak adjusted) analyses that are significant transcriptome-wide with both the obesity and the BMI tests. This includes hsa-miR-23b-3p (from female, obesity tests, adjusted p: 0.007 and 0.011, log FC: −0.344 and −0.240 from DESeq2 and LIMMA, respectively. From BMI LIMMA, adjusted p: 0.024 and 0.026, log FC: −0.016 and −0.014, from female and both sexes, respectively), which through Gene Ontology associations is involved with cardiac muscle cell growth and vascular permeability and growth, hsa-miR-15b-5p (female, obesity DESeq2 adjusted p: 0.014, log FC: −297, both sexes, BMI LIMMA adjusted p: 0.029, log FC: −0.010), associated with cardiac muscle hypertrophy and angiogenesis, and hsa-miR-96-5p (from both sexes, BMI LIMMA adjusted p: 0.043, log FC: −0.021, and from both sexes, obesity BMI test, adjusted p: 0.011 and 0.008, log FC: −0.788 and −0.692 from RIN3/BraakIV and RIN3/Braak adjusted, respectively), associated with vascular smooth muscle cell proliferation with cellular response to cholesterol. An additional 37 miRNA are significantly associated with BMI in the nucleus accumbens, both sexes (filtering RIN3/Braak adjusted). Among them are hsa-miR-25-3p (adjusted p: 0.037, log FC: −0.012) and from the obesity-implicated set, hsa-let-7b-5p (adjusted p: 0.047, log FC: −0.007), with associations for cardiac muscle tissue growth and for angiogenesis, respectively. The small RNA DESeq2 results are shown in Table 4 and Supplemental File 1 (ObesityBMI_Significant_Results.xls), second tab.

### Gene Set Enrichment Analysis of Obesity

Gene set enrichment analysis results for the filter RIN3/BraakIV are summarized by category in Table 5. Full obesity GSEA results are given in Supplemental File 3 (GSEA_Obesity_DESeq2.xls), with filtering and stratifications in separate tabs, and GSEA result categorization counts and ratios per analysis (in separate columns) in Supplementary File 4 (GSEA_DESeq2_categorizations.xls). The number of gene set results for each differential expression analysis ranges from 23 to 343. The most common categorization of the gene-sets based on key-word association is immune system or inflammation related. For example, the top three results from the hypothalamus, both sexes (filter RIN3/Braak4) are Reactome’s “cytokine signaling in immune system” and “innate immune system” and Kegg’s “cytokine-cytokine receptor interaction.” The most common categorizations for the hypothalamus/male, hypothalamus/both sexes and nucleus accumbens/male sets (filtering RIN3/BraakIV) are by far immune/inflammation related, with 132/300, 65/143 and 21/45 gene sets, respectively.

**Table 5:** Number of Gene Set Enrichment Analysis results from obesity differential expression results with DESeq2, in each brain region and sex stratification (RIN3/Braak4 sets), as categorized by gene-set name key-word association. Note that a gene set may have multiple categories, so the number of gene sets over all categories (first row) does not necessarily equal the sum of all other rows.

|  | All gene sets | Hypo/ M | Hypo/ F | Hypo/ Both | NucAcc/ M | NucAcc/ F | NucAcc/ Both |
| --- | --- | --- | --- | --- | --- | --- | --- |
| All categories | 1329 | 300 | 58 | 143 | 45 | 23 | 30 |
| None | 440 | 0 | 0 | 0 | 0 | 0 | 0 |
| Other | 214 | 53 | 25 | 34 | 11 | 2 | 10 |
| Obesity | 78 | 5 | 7 | 7 | 0 | 0 | 1 |
| Immune/Inflam | 229 | 132 | 6 | 65 | 21 | 8 | 0 |
| Neurons | 114 | 39 | 16 | 6 | 1 | 2 | 11 |
| Vascular system | 25 | 8 | 4 | 5 | 0 | 1 | 1 |
| Apoptosis | 39 | 22 | 0 | 4 | 4 | 1 | 0 |
| Cancer | 55 | 19 | 0 | 5 | 4 | 0 | 1 |
| Cell Cycle | 37 | 10 | 0 | 1 | 1 | 2 | 1 |
| DNA damage/repair | 16 | 3 | 0 | 0 | 1 | 0 | 1 |
| Development | 45 | 6 | 0 | 1 | 2 | 0 | 1 |
| Microtubules | 41 | 4 | 1 | 3 | 0 | 0 | 0 |
| TF targets | 32 | 13 | 0 | 6 | 2 | 0 | 0 |
| Transc/Transl | 89 | 22 | 2 | 17 | 1 | 8 | 4 |

\*Hypo: hypothalamus, NucAcc: nucleus accumbens, M: male, F: female, Immune: immune system,
Inflam: inflammation, TF: transcription factor, Transc: transcription, Transl: translation

Not surprisingly, many gene sets are neuron-related, the most common categorization for the nucleus accumbens/both sexes results (filtering RIN3/BraakIV). Four of the six gene sets among the filtering RIN3/BraakIV, have the Reactome category “Neuronal System” among the top results. Interestingly, the most significant result from the nucleus accumbens, both sexes (filter RIN3/Braak4) is Reactome’s “opioid signaling”, with adjusted p-value of 0.010. There are some obesity-related gene sets among the significant results, for example from the hypothalamus, female (filtering RIN3/Braak4) Kegg’s “insulin signaling pathway” with adjusted p-value of 0.015, but overall, obesity-related pathways are sparse.

### Brain Region Differences

Differential expression results between the hypothalamus and nucleus accumbens are abundant. There are many thousands of genes, and many hundreds of miRNAs that are significantly differentially expressed with adjusted p-value < 0.05 for each combination of cases/controls/both and male/female/both and for both DESeq2 and LIMMA with large overlap across them. There are more than 2,000 genes and nearly 900 miRNAs that are DESeq2 differentially expressed with the largest set of samples, both statuses and sexes combined (filtering RIN3/Braak adjusted). Of the top 10 most significant results there is very large overlap of genes and miRNAs across all sets, regardless of obesity status or sex stratification. Among these are some brain-related genes including dopamine receptor D1 and hippocalcin, and several miRNAs from the obesity-implicated set including hsa-miR-10b-5p and hsa-miR-539-3p, which are among the obesity/BMI differential expression results. Gene set enrichment results of the brain region DESeq2 results are also abundant. There are at least 150 gene-sets significantly associated with the rank of differential expression results for each analysis. Reactome’s “Cholesterol biosynthesis” is within the top 10 most significant results in each of the RIN3-filtered analyses, followed only by KEGG’s “Ribosome” and Reactome’s “Peptide chain elongation”. The top brain region DESeq2 result (top 10 most significant by p-value from each filtering and stratification analysis) are shown in Supplemental File 5 (Brain_region_DESeq2_Top10_Significant_Results.xls), mRNA and miRNA in separate tabs. GSEA results from brain region DESeq2 analyses are shown in Supplemental File 6 (GSEA_BrainRegion_DESeq2.xls), and gene set categorization information among the columns of Supplementary File 4 (GSEA_DESeq2_categorizations.xls).

### Sex Differences

Differential expression by sex yields at least a dozen significant mRNA results for the filter RIN3/BraakIV, with many hundreds from the largest stratification set, nucleus accumbens of all obesity statuses. Similar numbers were seen with other filter sets. This sample set also yields the most miRNA results, with more than a dozen for each test and more than 50 in most. This is compared to all other sample stratifications which, with one exception, have fewer than 4 significant results in all sample sets and tests. For both mRNA and miRNA there is large overlap between the DESeq2 and LIMMA results. Unsurprisingly, most of the top results are on the X and Y chromosomes. From the top ten most significant results of all RIN3/BraakIV DESeq2 analyses (stratifications of cases status and brain region), there are twenty-two mRNAs, none of which are autosomal. By expanding to the top fifty most significant results there are 297 mRNAs over all stratifications, of which 239 are autosomal. Of those 239, there are 9 from both brain regions in at least one analysis, including four immune-related genes (IGKC, IGHG2, ISG15 and NLRP2) and two lncRNAs. There are many mRNAs that appear among the top results in multiple analyses but only of one brain region, and similarly, a number from only obese or only control analyses (or combined), but never together. This is true for both large and small RNA analyses. There is an abundance of obesity-implicated miRNA within these results. Among the 454 unique miRNAs that appear within the top 50 results of any DESeq2 analysis, 53 are within our obesity-implicated set, 33 of which have negative direction of effect (higher in males), 12 positive, and 8 that differ depending on the context. Significant sex DESeq2 results are shown in Supplemental File 7 (Sex_Significant_Results.xls), mRNA and miRNA in separate tabs.

With the exception of nucleus accumbens, cases (filtering RIN3/BraakIV), each set of large RNA differential expression analyses yields Gene Set Enrichment Analysis results, shown in Supplemental File 8 (GSEA_Sex_DESeq2.xls), with resulting categorization information among the columns of Supplementary File 4 (GSEA_DESeq2_categorizations.xls). These GSEA results are heavily skewed towards those gene sets characterized as related to the immune system or inflammation, even more so than the obesity results and brain region GSEA results. For example, in analyses of both regions the gene set “autoimmune thyroid disease” from KEGG, categorized as immune-related and is also obesity-related, is significantly associated with the sex DE results (filtering RIN3/BraakIV) among all samples and among only obese samples.

### Validation

Brain region RNA sequencing DE results were successfully validated with significance (adjusted p-value < 0.05) and the same direction of effect with qPCR ΔΔCt analyses for all six transcripts in nearly all sample stratifications. Each of the chosen transcripts (ARP21, SYNDIG1L, DRD1, hsa-miR-139-5p, hsa-miR-552-5p and hsa-miR-10b-5p) had very significant RNA sequencing DE results in each of the sample stratifications used for the validation (female, RIN3/BraakIV and RIN6/BraakIV), with adjusted p-values ranging from 1.187E-05 to 2.313E-136. These DE results are validated by the qPCR data, with ΔΔCt analysis adjusted p-values ranging from 0.000548 to 2.058E-29. The only exceptions are among the RIN6/BraakIV, control, female sample stratifications, for which hsa-miR-139-5p and hsa-miR-10b-5p yielded directions of effect matching the RNA sequencing DE results and whose nominal p-values were significant but adjusted p-values were not (0.116 and 0.223). Obesity RNA sequencing differential expression validation is summarized in Table 6. The USP6 RNA sequencing DE result was validated by the ΔΔCt analysis with significance (adjusted p < 0.05) and fold change in the same direction. TTN and NDF ΔΔCt results also have fold change directions matching the RNA sequencing results and although the p-values are nominally significant, they are not significant after multiple hypothesis testing adjustment.

**Table 6:** Summary of the RNA sequencing differential expression results (adjusted p-value and mean ΔΔCt) and qPCR ΔΔCt results (adjusted p-value and RQ value) for the three transcripts chosen to validate obesity differential expression.

| Transcript name | Test | Stratification (all female) | DESeq2 adjusted p | $\Delta\Delta\text{Ct}$ adjusted p | DESeq2 log2 fold-change | Mean $\Delta\Delta\text{Ct}$ |
| --- | --- | --- | --- | --- | --- | --- |
| USP6 | Obesity | Hypothalamus, RIN6/BraakIV | 9.769E-07 | 0.026 | 1.781 | 8.687 |
| TTN | Obesity | Hypothalamus, RIN3/BraakIV | 0.025 | 0.222 | 0.725 | 0.975 |
| NDNF | Obesity | Hypothalamus, RIN3/BraakIV | 0.027 | 0.073 | 0.990 | 1.081 |

## Discussion

We present an analysis of large and small RNA sequencing in 444 samples after filtering for RIN and Braak stage to remove samples with degraded RNA and evidence for Alzheimer pathology. Obesity has not been previously studied with RNA sequencing in the human brain, and these samples give a rare opportunity to analyze the role of the hypothalamus and nucleus accumbens in obesity and BMI. Because of the neuroendocrine role of the hypothalamus and the reward/addiction role of the nucleus accumbens, each brain region has functions that are associated with obesity but we have for the first time evaluated their gene and miRNA expression association with the obese phenotype. We have shown that a small number of genes and miRNAs have statistically significant differential expression of obesity and BMI in these tissues, including some involved in metabolism such as APOBR and CES1, and a number associated with the immune system and inflammation, which is reinforced by gene set enrichment analysis.

Our criteria for samples’ case and control definitions give a binary contrast of stark phenotypic differences, requiring that the last four measures of BMI before death are greater than 30 or between 18.5 and 25. This is a conservative approach that removed from study individuals whose weight fluctuated or lost weight to the point of changing obesity status prior to death. In addition to this test of binary obese status with DESeq2 we conducted a Limma linear regression test with the continuous BMI measure, which included many subjects who did not meet the strict case/control criteria (see Table 1 for sample sizes of these analyses).

For test of obesity differential expression we used both DESeq2 and Limma linear regression. The implementation of Limma was intended to more thoroughly address differences observed between our two phases of sequencing. As DESeq2 cannot utilize sequencing counts that have been transformed, we accounted for sequencing phase among our potential DESeq2 model covariates, but with Limma we also used a data transformation with ComBat batch correction [55]. Ultimately the results of the two analyses were similar, indicating that the two methods for accounting for phase differences may be comparable.

A small number of differential expression results were statistically significant. This may indicate that there are few genetic signals of obesity to be detected in the hypothalamus or nucleus accumbens or that such effects are small and that more samples providing additional power are needed for detection. While the total number of samples studied here is relatively large, it is compromised by the intrinsic challenges of using human post-mortem samples from these longitudinal studies of aging adults, as many were removed for low RNA integrity and high Braak stage. Inherent with human samples is large heterogeneity which will obfuscate true associations, and inherent with post-mortem samples is the possibility that a true obesity signal is overwhelmed by death processes. However, these limitations do not outweigh the clear benefit of studying the disease in human tissue directly without an animal model of disease, or of the unique interrogation of brain tissue with regard to obesity and BMI. Our large sample size has at least partially overcome those inherent difficulties, as a handful of interesting genes and miRNAs are observed whose expressions are significantly associated with obesity status or BMI.

The RNA sequencing analyses were validated with ΔΔCt analyses from qPCR data in the same samples for the brain region DE of six transcripts and the obesity DE of three transcripts. Female stratified analyses were chosen, as representative of all analyses but with feasible sample size. Each RNA sequencing DE result had power for validation above 0.95, based on sample size and DESeq2 effect size and p-value. Although performed in a limited number of transcripts, this validates that the RNA sequencing data is comparable to qPCR data.

Inflammation and the immune system are the among the most prominent signals seen in our obesity differential expression and gene set enrichment results, in the hypothalamus by genes such as SLAMF8, C1QTNF4, MUC16, FCGBP and CXCL8 and 65 of the 143 significant gene sets from the analysis of both sexes. The analysis of the nucleus accumbens has far fewer inflammation mRNA results, but has far more significant miRNAs with vascular associations, such as angiogenesis, a symptom of chronic inflammation. These include hsa-miR-210-3p, hsa-miR-23b-3p, hsa-miR-15b-5p, hsa-miR-96-5p,hsa-miR-96-5p and hsa-let-7b-5.

Tissue inflammation has been associated with obesity, not only the system-wide inflammation observed in many organ systems, but also specifically in the hypothalamus and some other brain tissues [56]. There is evidence that inflammation of the hypothalamus precedes that of peripheral tissues [57], suggesting that inflammation-induced dysregulation of the energy-regulating functions of the hypothalamus contribute to comorbidities such as insulin resistance, and to maintaining over-eating and obesity [58, 59]. Although this may suggest that the signals of hypothalamic inflammation we have ascertained from gene expression may be partially causal, it would be difficult to disentangle the two potential types of inflammation, causal and resultant, because the obese individuals in this study may have experienced generalized inflammation of brain tissue for many years. The differences in the inflammation signal between the hypothalamus and nucleus accumbens may indicate otherwise, but we should also be cautious that the inflammation signals may be due to technical artifacts. For example, because adjacent brain regions may have great differences in vascular characteristics, signals of vascular differences may be particularly susceptible to slight differences in tissue composition due to dissections. In addition, inflammation-associated results may generally be more likely to be a result of type I error, due to a characterized bias towards immune-related genes and pathway over-representation in differential expression analyses regardless of biological context [60].

For example, because adjacent brain regions may have great differences in vascular characteristics, signals of vascular differences may be particularly susceptible to slight differences in tissue composition due to dissections. In addition, inflammation-associated results may generally be more likely to be a result of type I error, due to a characterized bias towards immune-related genes and pathway over-representation in differential expression analyses regardless of biological context [60].

Carboxylesterase 1 (CES1) and Apolipoprotein B Receptor (APOBR) are both DESeq2 genomewide significant in at least one subset of hypothalamus samples. Expression of CES1 has been previously associated with obesity and type 2 diabetes in human adipose tissue [51]. It plays a role in both toxin, including cocaine and heroin, detoxification but also in cholesterol and lipid metabolism. The nucleus accumbens is involved with drug addiction and impulse control, but CES1 was genome-wide significant only in the hypothalamus. Given the association of CES1 with obesity in adipose tissue and now in the hypothalamus, it is possible that the mechanism of this association is driven by the role of CES1 in the metabolism of cholesterol and lipids rather than toxins.

APOBR is more lowly expressed in our obese samples relative to control in the hypothalamus, for both sexes. It binds APOB, mediating uptake of lipids into the cell. Lower expression of this receptor may cause lipid deficits and negatively impact cellular health and function. The effect of lipid metabolism on brain and neuron health has been thoroughly analyzed especially with regard Alzheimer’s disease and APOE, but also APOB [61]. Dysregulation of the lipid intake may negatively affect the hypothalamus’ normal functions regulating energy intake and expenditure, potentially helping to explain the observed association. Alternatively, reduced ability to respond to APOB may be interpreted by the hypothalamus as a signal from other tissues, disrupting normal hypothalamus regulation. Although there are a handful of obesity associated GSEA results from both the nucleus accumbens and hypothalamus, most GSEA results are immune system, inflammation or nervous system related. It is interesting, however, that the gene set most significantly associated with the nucleus accumbens, both sexes DESeq2 results is “opioid signaling,” a set of 72 genes. This may indicate that the nucleus accumbens’ role in regulating impulse control and addiction may indeed also play a role with obesity, as was the original rationale to include nucleus accumbens in this study.

In order to interrogate possible interaction between obesity and brain region, differential expression of the hypothalamus and nucleus accumbens was conducted. These analyses yield a very large number of significantly differentially expressed genes and miRNAs. Gene set enrichment analysis from the mRNA DESeq2 are also numerous, but include some that are potentially obesity associated, such as “Cholesterol biosynthesis”. In addition, the miRNAs that are significantly differentially expressed by brain region include several that are among the obesity/BMI differential expression results. These associations with obesity may be unsurprising given the two very different functions of the hypothalamus and nucleus accumbens and their potential roles in obesity. Similarly, the miRNAs that are significantly differentially expressed by sex include many within our *a priori* defined obesity-implicated set, perhaps showing an interaction of obesity and sex in these tissues and justifying stratified analyses. GSEA with the DESeq2 tests of sex differential expression results yield many gene sets characterized as related to the immune system or inflammation, even more than from the analyses of obesity status and brain region, perhaps indicating sex-differences in the obesity-associated inflammation of the brain tissue or of the tissue’s ability to maintain its functions while experiencing inflammation.

Differential expression of obesity and BMI yielded few significant results, but particularly notable are the mRNAs ABOR and CES1 which linked to metabolism and obesity, as well as a number of other mRNAs and miRNAs showing aspects of brain inflammation. The gene set enrichment results reiterate this signal of inflammation and immune system differences by obesity status, in addition to angiogenesis. In addition, “Opioid signaling” is among the gene sets associated with the nucleus accumbens obesity differential expression results. If the per-transcript signal of obesity in these tissues is difficult to detect, it appears that there is more information captured on the level of gene sets, reinforcing that there are legitimate patterns of obesity taking place in the hypothalamus and nucleus accumbens. The analyses of differences by brain region show huge differences between the hypothalamus and nucleus accumbens with an indication of a BMI association from the GSEA results, and like the obesity analysis, analysis of sex indicates differences of the immune system and inflammation.

## Conclusions

We have conducted differential expression analyses of obesity and BMI in humans with RNA sequencing of 207 hypothalamus and 276 nucleus accumbens post-mortem samples from the Framingham Heart Study, Religious Orders Study and Memory Aging Project. These analyses were conducted both transcriptome-wide and with sets of mRNAs and miRNAs that were *a priori* defined to be associated with obesity or BMI. Our study may be under-powered to detect transcript-level results, despite our relatively large sample size. With significance of adjusted p-value less than 0.05 there are sixteen mRNAs and five miRNAs with significant differential expression by obesity or BMI over all analyses and RIN and Braak filtering criteria. Among the obesity DE mRNAs are CES1, associated with detoxification and cholesterol and lipid metabolism, and APOBR, a receptor for APOB which mediates cellular lipid uptake. Many of the significant miRNA differential expression analyses are associated with the immune system, angiogenesis and inflammation, signals which are reiterated by many gene set enrichment analysis results from the mRNA differential expression results. Despite few transcript-level results, there appear to be valid signals of obesity and BMI detected in these tissues at the level of genesets, dominated by signals of inflammation.

## Data Availability

Data for ROSMAP can be requested on the RADC Resource Sharing Hub at www.radc.rush.edu. Data for FHS is in the process of being submitted to dbGAP (accession number phs002611.v1.p1).

## Acknowledgements

We would like to acknowledge the Boston University Bioinformatics Program.

## Supplementary Materials

Supplementary File 1 is an excel file which contains the large and small RNA results (in separate tabs) of the obesity and BMI tests from all sample sets. For each transcript with a significant result (rows), the transcript ID and name, chromosome, and a brief description are included (columns), as well as the sample subset for which the transcript was significant (the same transcript in different sample subsets are in separate rows), as well as DESeq2 information (log fold-change, base mean, nominal p-value, adjusted-pvalue and obesity-set adjusted p-value) and LIMMA information (log fold-change, nominal p-value, adjusted p-value and obesity-set adjusted p-value).

Supplementary File 2 is an excel file which summarizes the number of results from all obesity, BMI, brain region and sex analyses. Separate analyses (mRNA/miRNA and sample filter) are shown in separate tabs, e.g. “mRNA_Filter_RIN3Braak4”. Within each tab, rows distinguish the test (e.g. obesity status) and sample set (e.g. hypothalamus/male), and further columns show the sample sizes, covariates included in the analysis, the number of significant DESeq2 results, number of significant LIMMA results, and the number that overlap between the two.

Supplementary File 3 is an excel file containing the GSEA results from the obesity DESeq2 analyses in totality. Different analyses (stratifications and sample filtering) are displayed in separate tabs. In addition to the GSEA outputs, each result (row) also shows the key-word categorization of the pathway, and a list of other analyses in which the pathway was significant result.

Supplementary File 4 is an excel file which contains GSEA result categorization information, counts and ratios of counts in separate tabs. For each separate analysis (columns) and for each combination of gene set categorization (rows), the count (number of GSEA results) and ratio of counts (count divided by the total number of GSEA results) is shown.

Supplemental File 5 is an excel file which shows the top brain region DESeq2 result (top 10 most significant by p-value from each filtering and stratification analysis), mRNA and miRNA in separate tabs. Each row is a transcript and among the column information are the gene name, chromosome, brief description, the specific analyses (and number) in which the transcript was among the top results, adjusted p-values for separate analyses, and comma-delimited per cell in the same order as the analyses listed, the log fold-changes, DESeq baseMean values, and sample sizes.

Supplemental File 6 is an excel file containing the GSEA results from brain region DESeq2 analyses. Different analyses (stratifications and sample filtering) are displayed in separate tabs. In addition to the GSEA outputs, each result (row) also shows the key-word categorization of the pathway, and a list of other analyses in which the pathway was significant result.

Supplemental File 7 is an excel file which shows the results from sex DESeq2 analyses, mRNA and miRNA in separate tabs. For each transcript with a significant result (rows), the transcript ID and name, chromosome, and a brief description are included (columns), as well as the sample subset for which the transcript was significant (the same transcript in different sample subsets are in separate rows with ‘-integer’ appended to the ensemble ID row names), as well as DESeq2 information (log foldchange, base mean, nominal p-value, adjusted-pvalue and obesity-set adjusted p-value) and LIMMA information (log fold-change, nominal p-value, adjusted p-value and obesity-set adjusted p-value).

Supplemental File 8 is an excel file containing the GSEA results from sex DESeq2 analyses. Different analyses (stratifications and sample filtering) are displayed in separate tabs. In addition to the GSEA outputs, each result (row) also shows the key-word categorization of the pathway, and a list of other analyses in which the pathway was significant result.

## Author Contributions

The study was conceived and designed by ALD, RHM, and SS with input from DAB. Brain dissections were performed by TS, with input from AM, and JAS with additional lab work performed by CW, RHM, and JB. CW performed the data analyses with input from AL, ALD and RHM, and drafted the manuscript, with critical review from all authors.

